# IL-1α secreted by subcapsular sinus macrophages promotes melanoma metastasis in the sentinel lymph node by upregulating STAT3 signaling in the tumor

**DOI:** 10.1101/2021.10.11.463954

**Authors:** Tommaso Virgilio, Joy Bordini, Giulio Sartori, Irene Latino, Daniel Molina-Romero, Cristina Leoni, Murodzhon Akhmedov, Andrea Rinaldi, Alberto J. Arribas, Diego Morone, S. Morteza Seyed Jafari, Marina Bersudsky, Aner Ottolenghi, Ivo Kwee, Anna Maria Chiaravalli, Fausto Sessa, Robert E. Hunger, Antonino Bruno, Lorenzo Mortara, Elena Voronov, Silvia Monticelli, Ron N. Apte, Francesco Bertoni, Santiago F. Gonzalez

## Abstract

During melanoma metastasization, tumor cells originated in the skin migrate via lymphatic vessels to the sentinel lymph node (sLN) in a process that facilitates their spread across the body. Here, we characterized the innate inflammatory responses to melanoma metastasis in the sLN. For this purpose, we confirmed the migration of fluorescent metastatic melanoma cells to the sLN and we characterized the inflammatory response in the metastatic microenvironment. We found that macrophages located in the subcapsular sinus (SSM), produce pro-tumoral IL-1α after recognition of tumor antigens. Moreover, we confirmed that the administration of an anti-IL-1α depleting antibody reduced metastasis. Conversely, the administration of recombinant IL-1α accelerated the lymphatic spreading of the tumor. Additionally, the elimination of the macrophages significantly reduced the progression of the metastatic spread. To understand the mechanism of action of IL-1α in the context of the lymph node microenvironment, we applied single-cell RNA sequencing to dissected metastases obtained from animals treated with an anti-IL-1α blocking antibody. Amongst the different pathways affected, we identified STAT3 as one of the main targets of IL-1α signaling in metastatic cells. Moreover, we found that the anti-IL-1α anti-tumoral effect was not mediated by lymphocytes, as IL-1R1 KO mice did not show any improvement in metastasis growth. Finally, we found a synergistic anti-metastatic effect of the combination of IL-1α blocking and the STAT3 inhibitor (STAT3i) stattic. In summary, we described a new mechanism by which SSM support melanoma metastasis, highlighting a new target for immunotherapy.

## INTRODUCTION

Melanoma is the most lethal form of skin cancer and a serious threat for public health. In recent years, the incidence of this type of cancer has progressively increased and it is currently one of the most common malignancies in both adult and young individuals^1^. During melanoma development, malignant cells in the skin acquire genetic mutations that lead them towards the lymphatic vessels, which serve as a transportation system^2^. Once in the lymphatics, the metastatic cells initiate an active migration that leads them towards the sentinel LN (sLN)^3^. The presence of melanoma metastasis in this organ is indicative of a poor prognosis, drastically decreasing the survival rate of the patients^4,5^.

Upon breaching the LN capsule, metastatic cells access the LN sinuses via the afferent lymphatics, following chemokine gradients generated by lymphatic endothelial cells^6–8^. The invasion of the sLN is initiated in the subcapsular sinus area (SS)^6,8,9^ and progressively spreads towards the inner structures of the sLN^10^. This process facilitates the access of the metastatic cells to the bloodstream via high-endothelial venules (HEVs) and their consequent spread to distant organs^11–13^.

The LN sinuses are populated by resident phagocytic cells, including three distinct macrophage subsets called subcapsular sinus macrophages (SSM), medullary macrophages (MM) and medullary cord macrophages (MCM), according to the area they reside^14^. Strategically positioned along the SS area, SSM are the first immune cells to encounter lymph-transported antigens and pathogens, preventing their systemic dissemination^15^. In addition, they play a critical role in the initiation of the immune responses against immune complexes and virus^16–18^ as well as promoting humoral immunity^19,20^. Despite the role of macrophages against infectious pathogens has been largely demonstrated, their involvement in the response against tumor remains somehow controversial^21–27^. This is mainly due to the ability of these cells to activate either anti- or pro-tumoral responses, according to their cell plasticity, that allows them being dicotomically classified in M1 and M2 macrophages^28,29^. For instance, some authors have described a protective function of SSM, which was associated with the capturing of dead tumor cell-derived antigens^23^ and their cross-presentation to CD8^+^ T cells^21^. Moreover, Tacconi and colleagues have recently suggested a protective role of CD169^+^ LN macrophages in breast cancer metastasis, which was dependent on the presence of B cells^27^. Conversely, other studies have revealed a pro-tumoral effect of these cells, mainly linked with their capacity to trigger and maintain the inflammatory response both in peripheral and lymphoid tissues^9,30,31^.

The inflammatory response plays a fundamental role in the behavior of cancer cells. Some cancers, including melanoma, are able indeed to grow in chronically inflamed conditions and to take advantage of inflammation^32–35^. One of the mechanisms by which innate inflammation supports tumor growth is by the release of IL-1 family cytokines^36,37^. IL-1β, the major component of this family, has been shown to endorse different tumors, mainly by mediating immune suppression and by activating endothelial cells^38,39^. Indeed, recent evidence suggests that blocking the IL-1R signaling might prolong the survival time of patients with different tumors^40–44^. In addition, IL-1β antagonism can synergize with immune checkpoint inhibitor therapy^38^. However, the mechanisms responsible for this process might vary between the primary tumor and the metastatic areas, including the sLN^45,46^. Understanding these differences will influence the design of specific immunotherapies intended to control tumor dissemination in both locations^47,48^ and in different types of tumors, including melanoma^49,50^.

In the present work we characterized the innate immune response of the sLN to melanoma metastasis invasion. Furthermore, we identified a novel mechanism that associates the inflammatory reaction, initiated by SSM, to the progression of the metastatic melanoma cells. These results will have a potential impact generating new therapies and improving the efficiency of the current immunotherapies against metastatic melanoma, possibly acting on macrophages, that represent the most abundant inflammatory cells infiltrating tumor.

## RESULTS

### Development and characterization of a murine model of melanoma metastasis in the draining popliteal lymph node

To study the metastatic process in the sentinel lymph node (sLN) we transduced the melanoma cell line B16-F1 with a lentiviral vector codifying for mCherry and we characterized the expression of this fluorescent protein by FACS and microscopy (Fig. 1A and Supp. Fig. 1A, respectively). The primary tumor was induced by subcutaneous injection of the cancer cells in the mouse footpad, similarly to what was previously reported (Fig. 1B)^51^. Next, the formation of the primary tumor was monitored by measuring tumor volume (Supp. Fig. 1B,C) and tumor fluorescence was quantified by using the *In Vivo* Imaging System (IVIS; Fig. 1C,D). Following this approach, we observed a significant engraftment starting from week one post tumor implantation (p.t.i) (Fig. 1D). Subsequently, we identified macrometastases in the sLN at three weeks p.t.i. (Fig. 1E). To study in more detail the progression of melanoma cells towards the sLN, we employed flow cytometry, observing a significant increase of the metastatic cells at day 15 p.t.i. compared to control samples (Fig 1F). In addition, to characterize the area of the metastasis, we used confocal microscopy analysis of the sLN at different time points following tumor induction (Fig. 1G). We measured a significant increase in the metastatic area starting at two weeks (Fig. 1H). Besides, the morphology of sLN metastases showed that, in concordance with previous works in humans^8^, metastatic cells initially invade the subcapsular sinus area (SS) (Fig. 1G,I). Furthermore, at later time points, the metastasis progressively expanded through the interfollicular area (IF), invading the transverse sinus (Fig. 1I, Sup Fig. 1D). Conversely, we did not observe the presence of metastatic cells in distant organs, such as the spleen or the lungs, at equivalent time points (Fig. 1J), confirming the lymphatic dissemination of the tumor.

**Figure 1.**
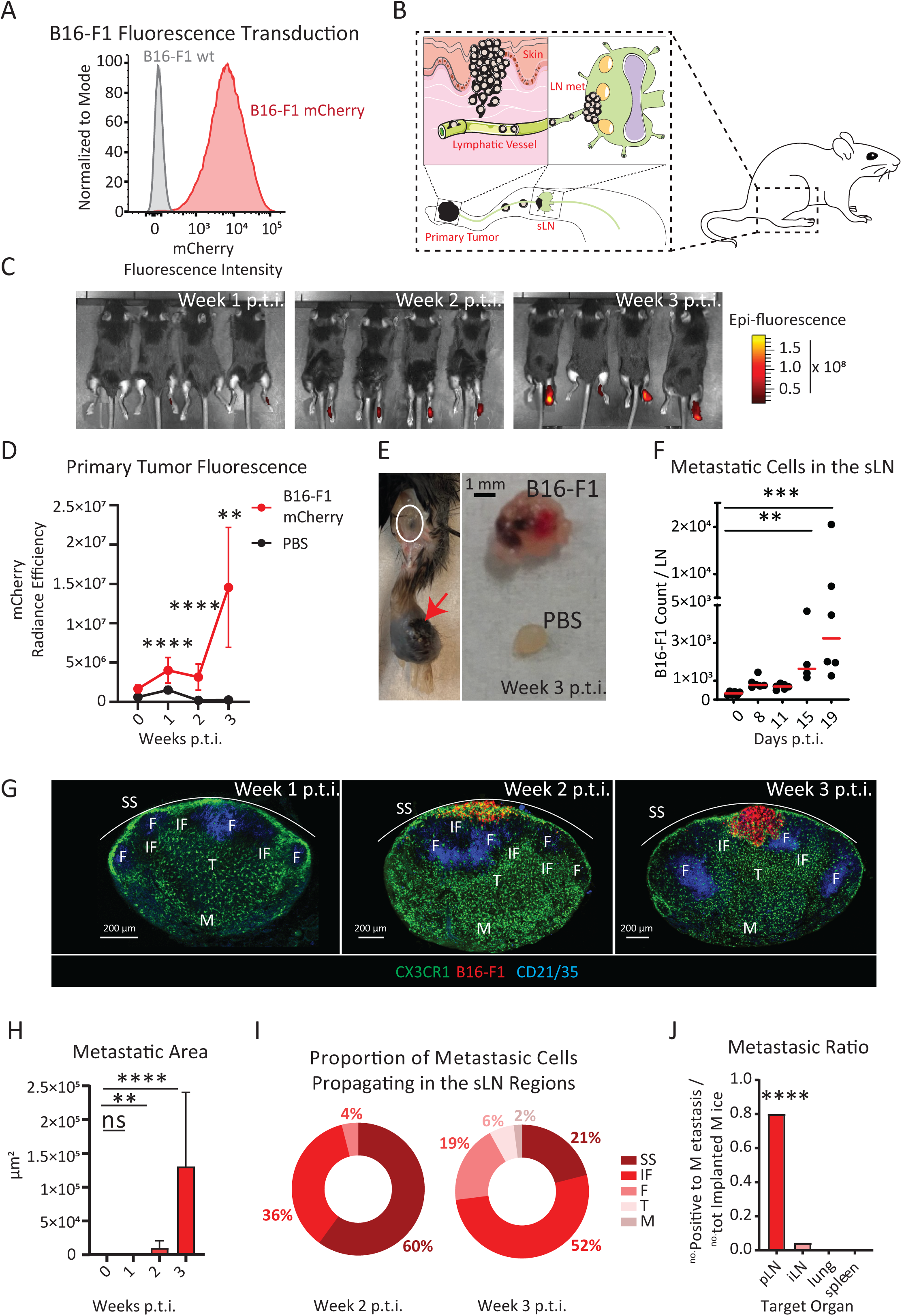
Mouse melanoma metastases growth in the sentinel LN. (A) FACS plot showing fluorescent expression of the mCherry transduced B16-F1 melanoma cells. (B) Schematic representation of the tumor model, including primary tumor engraftment (left) and migration of cells to the sLN (right). (C) Representative images and (D) quantification of IVIS time-course showing increasing primary tumor fluorescence (red). (E, left) Primary tumor (red arrow), draining sentinel LN (white circle) and (E, right) comparison between metastatic and healthy LN at week three p.t.i.. (F) Time-course of metastatic cell invasion of the sLN quantified by FACS. (G) Confocal micrograph of sLN at week one, two and three p.t.i., showing the position of B16 melanoma (red) with respect to CD21/35^+^ (blue) follicular dendritic cells and CX3CR1^+^ (green) myeloid cells. SS, IF, F, T and M stand for subcapsular sinus, inter-follicular, follicular, T cells and medullary areas, respectively. (H) Quantification of total metastatic area in the sLN, measured by confocal microscopy. (I) Quantification of tumor cells in the different compartments of the LN at week two and three p.t.i.. (J) Metastatic ratio, defined as the number of mice with metastases in the target organ divided by the total number of implanted mice, at week three p.t.i.. iLN stands for inguinal LN.

### IL-1α promotes melanoma growth in the sLN

To characterize the inflammatory reaction induced by metastasis development, we quantified the total number of immune cells infiltrated in the sLN, observing a significant two-fold increase starting from the first week p.t.i. (Fig. 2A). More in detail, increases in dendritic cells (MHC-II^+^, CD11c^high^, CD11b^+^ and CD11b^-^), NK cells (CD3^-^, NK1.1^+^), neutrophils (MHC-II^-^, GR1^high^), monocytes (MHC-II^-^, GR1^int^) and macrophages (MHC-II^+^, CD11c^intlow^, CD11b^+^, Supp. Fig. 2A), as well as B (B220^+^) and T cells (CD4^+^, CD8^+^, and FOXP3^+^ T_reg_), were observed (Supp. Fig. 2B). To further characterize the recruitment of immune cells to the metastasized sLN, a multiplexed approach was applied to quantify the concentration of different inflammatory chemokines, including CXCL13, CXCL9, CCL22, CCL5 and CCL2, in the sLN supernatant, observing a significant increase at week three p.t.i. (Supp. Fig. 2C). Additionally, we measured the concentration of other 13 inflammatory cytokines. Among the molecules analyzed, IL-1α exhibited a significant increase at week three p.t.i., compared to the control group. (Fig. 2B). A further study of the dynamics of IL-1α release highlighted that the upregulation started at week two p.t.i. (Fig. 2C). To evaluate if IL-1α secretion was associated with other tumor models, including another solid tumor infiltrating the sLN, we measured the level of this cytokine in sLN metastasized with melanoma B16-F10, or with the breast cancer cell line E0771, observing similar levels of IL-1α in both models at three weeks p.t.i. (Fig. 2D).

**Figure 2.**
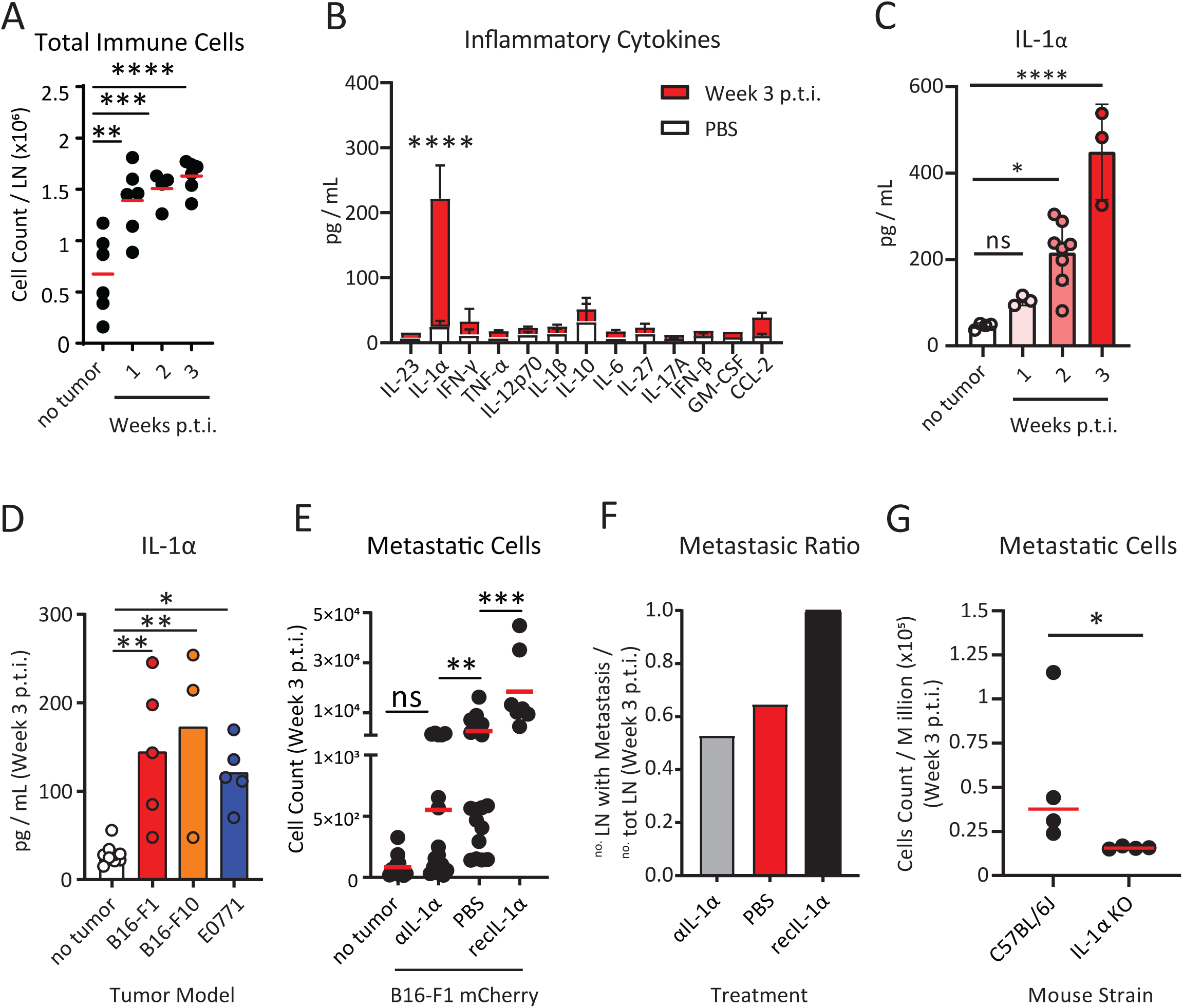
Pro-tumoral release of IL-1α in the metastatic LN. (A) Progressive increase in the size of the sLN correlated with the increase in the total number of immune cells, measured by flow cytometry. (B) Quantification of inflammatory cytokines in the supernatant of metastatic (red) and non-metastatic (white) LN. (C) Time course kinetics showing IL-1α release in the sLN during the first three weeks p.t.i.. (D) Quantification of IL-1α in the sLN at three weeks p.t.i. of different cancer models, including breast cancer (E0771) and the melanoma B16-F10. (E) Flow cytometric quantification of LN metastatic cells in animals treated with IL-1α depleting antibody or recombinant IL-1α in comparison to B16-F1 untreated group and PBS injected. (F) LN metastatic ratio in mice untreated or treated with anti-IL-1α antibody or recombinant IL-1α at week three p.t.i.. (G) Metastatic cells in the sLN of wild type and IL-1α KO mice three weeks p.t.i..

Next, we hypothesized a pro-tumoral role of IL-1α in the metastatic context. To explore this hypothesis, we treated mice with a daily subcutaneous injection of either IL-1α depleting antibody or recombinant IL-1α protein. Interestingly, blocking the IL-1α pathway by administration of the neutralizing antibody significantly decreased the metastasis growth in the sLN, as indicated by a reduction in the number of metastatic cells measured by flow cytometric analysis at week three p.t.i.. Conversely, the number of melanoma cells significantly increased in the sLN treated with recombinant IL-1α at equivalent time points (Fig. 2E). Moreover, the metastatic ratio, defined as the number of mice developing sLN metastasis at week three p.t.i. divided by the total number of mice showing primary tumor engraftment, was higher in the animals treated with recombinant IL-1α and it was reduced following IL-1α depletion (Fig. 2F). Nevertheless, the observed variation in the metastasis size after treatment could be dependent on the size of the primary tumor. Therefore, to discard that possibility, we normalized the number of metastatic cells to the primary tumor volume, confirming the results previously observed in Fig. 2E (Supp. Fig. 2D). Additionally, the previously described treatments did not have a significant effect on the growth of the primary tumor in comparison to the untreated control group (Supp. Fig. 2E). However, we observed that IL-1α KO mice showed a reduction not only of the metastatic cells at week three p.t.i. (Fig. 2G), but also of the primary tumor volume starting from the fourth week p.t.i. (Supp. Fig. 2F). This discrepancy could be partially explained by the mode of administration of the treatment, which promotes the transport towards the draining lymphatics, or by the time of administration of the compounds in comparison to the constant absence of IL-1α in the tumor microenviroment in KO mice.

### Subcapsular sinus macrophages are the main source of pro-tumoral IL-1α and disappear after tumor phagocytosis

In a previous study we characterized the role of LN macrophages as the main producers of IL-1α in the LN, following influenza vaccination^20^. To elucidate the main source of IL-1α in the melanoma model, we analyzed the infiltrating immune cells from the metastatic regions of the LN, by single cell RNA sequencing (scSeq, Fig. 3A). Following this approach, we confirmed that LN macrophages are the main producers of IL-1α (Fig. 3 B, C). Moreover, the depletion of this population, following the injection with clodronate liposomes (CLL), significantly reduced the levels of IL-1α in the LN supernatant (Fig 3D). Importantly, depletion of macrophages following CLL administration completely abrogated the growth of the metastatic melanoma cells in the LN, confirming their pro-tumoral nature (Supp. Fig. 3A, B). However, the local administration of CLL did not affect the volume of the primary tumor (Supp. 3C). Next, to identify the specific subset of macrophages responsible for the production of IL-1α, we used flow cytometry (Fig. 3E, F) and confocal microscopy (Fig. 3G), which pointed out SSM (CD169^+^, F4/80^-^) as the main source of IL-1α in the metastatic region. To prove the relevance of these findings also in humans, we performed immunohistochemical staining of melanoma metastatic LN from patients, confirming that the local production of IL-1α was associated with CD68^+^ tumor infiltrated macrophages in the SS region (Supp. Fig. 3D). As suggested by other studies^23^, we also reported that tumor infiltrating SSM phagocyte melanoma cells (Fig. 3G, H, Supp. Fig. 3E, Supp. Mov. 1).

**Figure 3.**
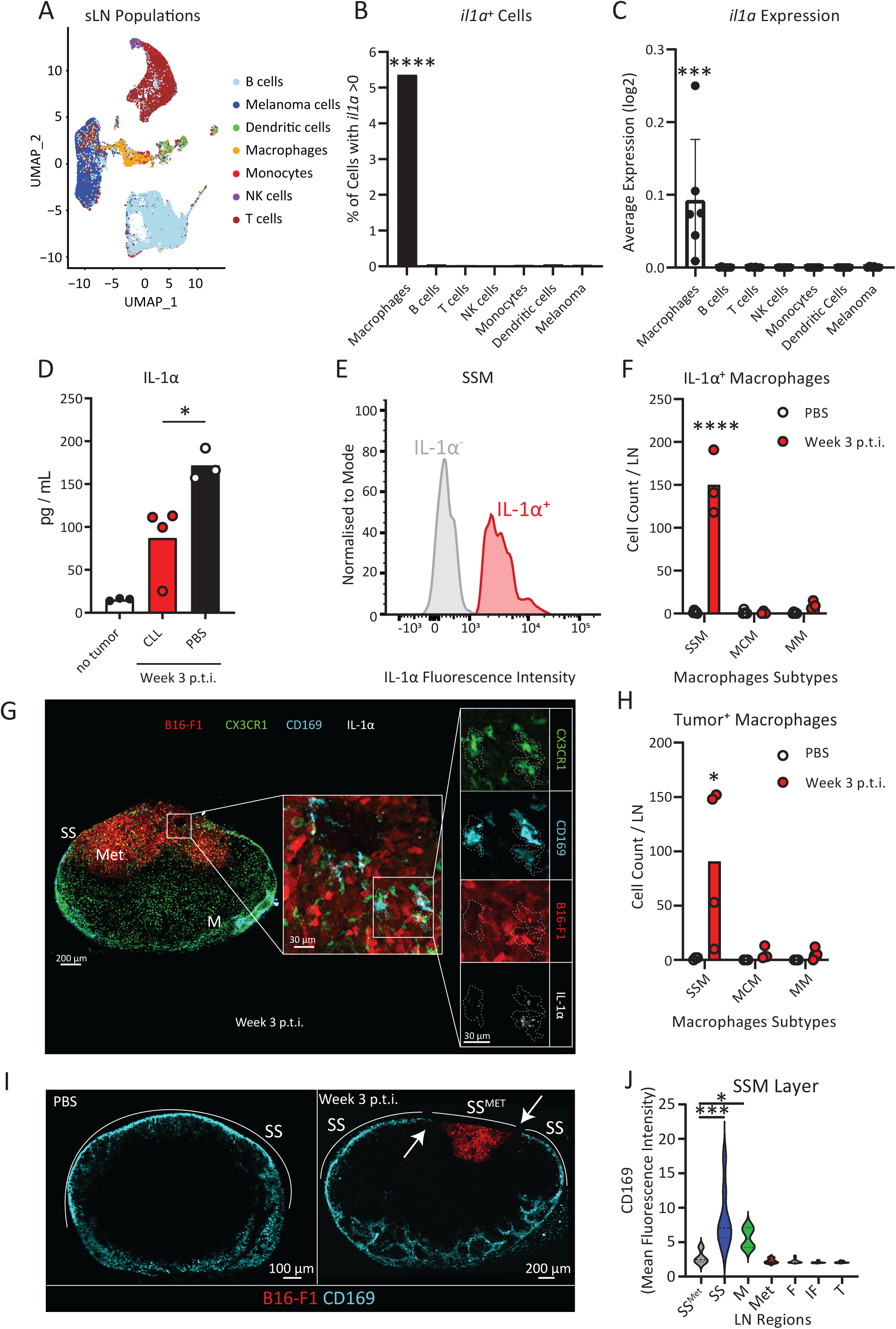
SSM are the main source of IL-1α. (A) UMAP plot of cell populations, identified by scSeq, in the metastasized sLN three weeks p.t.i.. (B) Percentage of cells expressing il1a and (C) average il1a expression in the cells of the sLN three weeks p.t.i., measured by scSeq. (D) IL-1α quantification in metastasized sLN supernatant of mice depleted for macrophages by clodronate liposome (CLL) injection in comparison to untreated metastasized and non-metastasized LN. (E) Flow cytometric histograms showing presence, three weeks p.t.i., of IL-1α^+^ (red) and IL-1α^-^ negative (gray) SSM. (F) Flow cytometric quantification of the number of IL-1α^+^ cells among the three major subtypes of macrophages in the sLN three weeks p.t.i. in comparison to negative controls. SSM, MCM and MM stand for Subcapsular Sinus Macrophages (CD169^+^ F4/80^-^), Medullary Cord Macrophages (CD169^-^ F4/80^+^) and Medullary Macrophages (CD169^+^ F4/80^+^), respectively. (G) Confocal micrograph showing the whole sLN (left) and magnifications of the metastatic region (center and right) indicating IL-1α And tumor vesicles in CX3CR1^+^ CD169^+^ macrophages. Colors indicate CX3XR1^+^ cells (green), mCherry^+^ melanoma (red), CD169^+^ macrophages (cyan) and IL-1α (white). (H) Flow cytometric quantification indicating the number of each subtype of tumor^+^ macrophages. (I) Confocal representative images of CD169^+^ macrophages distribution in the sLN three weeks p.t.i. in comparison to negative controls. (J) Quantification of CD169 fluorescence in the main regions of the LN three weeks p.t.i., indicating disruption of CD169 layer (white arrows) in the SS overlying the metastatic area (SS^MET^).

To investigate the mechanism of release of IL-1α by SSM, we quantified cell numbers by flow cytometry, observing that the total number of SSM remained constant during the first three weeks p.t.i. (Supp. Fig. 3F), while their frequency decreased (Supp. Fig. 3G). This was in contrast with a significant increase in the total number of macrophages observed at equivalent time points (Supp. Fig. 2A). Therefore, to investigate if SSM disappear following metastasis growth, we quantified by confocal microscopy the expression of the macrophage marker CD169 in different regions of the metastatic sLN. Interestingly, we observed that the CD169 layer in the SS was absent in the proximity of the metastatic area (Fig. 3I). More in detail, CD169 in the SS surrounding the metastatic region was expressed significantly less than in the non-metastasized SS (Fig. 3J), suggesting that SSM in direct contact with melanoma might undergo a cell death process. To confirm that the phagocytosis of tumor cell debris was able to induce SSM disappearance we injected B16-F1 lysate in the mouse footpad and performed flow cytometric analysis at 12 and 24 h following injection. We observed that the percentage of SSM significantly decreased compared with non-injected controls (Supp. Fig. 3H).

### SSM-derived IL-1α induces melanoma proliferation

In previous studies, we have demonstrated the involvement of IL-1α in the inflammatory reaction in the LN^20,52^. However, we did not observe here any significant effect in the recruitment of the major immune cells subtypes in the sLN following treatment with anti-IL-1α (Supp. Fig. 4 A, B). To further characterize the pro-tumoral mechanism of IL-1α, we measured the expression of IL-1R1, the only known receptor involved in the signaling of IL-1α^39^, in the infiltrated cell types of a metastatic sLN. Amongst the evaluated cells, NK cells and melanoma displayed the highest level of IL-1R1 expression (Fig. 4A). To clarify the role of immune cells in mediating the pro-tumoral function of IL-1α, we induced melanoma in IL-1R1 KO mice, in which tumor microenvironment cells can’t be involved in IL-1 signaling and only the tumor cells express this receptor. The absence of IL-1R1 in the immune compartment did not significantly affect neither the metastasis growth in the sLN (Fig. 4B) nor the metastatic ratio (Fig. 4C), demonstrating that the pro-tumoral effect of IL-1 signaling was not associated with the immune cell response. Next, after confirming the expression of IL-1R1 in cultured melanoma cells (Supp. Fig. 4C), we measured their proliferation rate following exposure to IL-1α. We discovered that IL-1α significantly promoted the proliferation of melanoma in murine and human cell lines (Fig. 4D and 4E, respectively). To further characterize the activation of IL-1R1 signaling in B16-F1 melanoma cells, we measured by qPCR the expression of the gene codifying for the Myeloid Differentiation Primary Response 88 protein (MYD88), the main mediator of the Interleukin-1 receptor associated kinase (IRAK) signaling activated by IL-1R1^39^. The results confirmed that melanoma cells treated with recombinant IL-1α significantly upregulated the *Myd88* gene (Fig. 4F).

**Figure 4.**
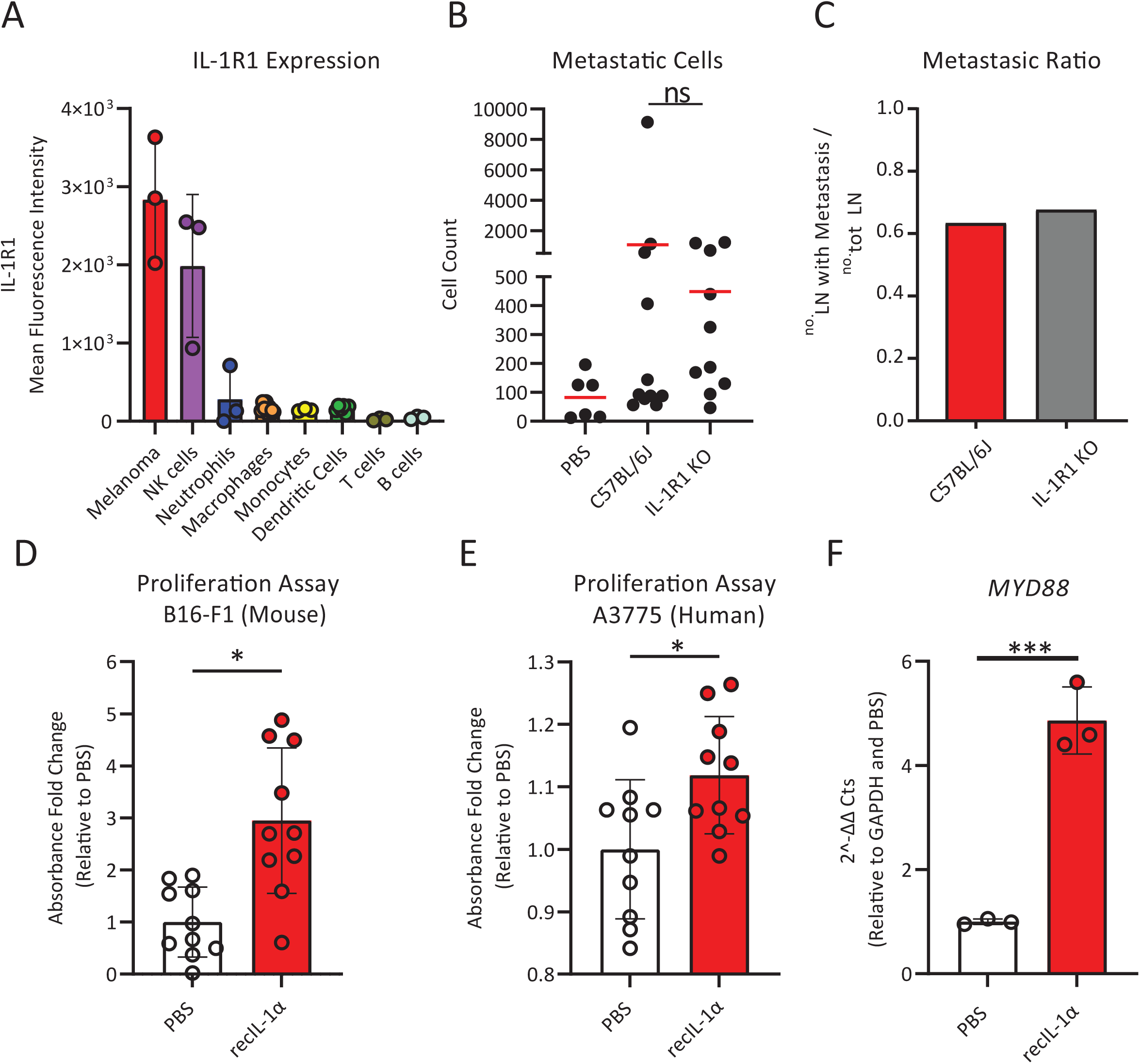
Direct effect of pro-tumoral IL-1α on metastatic cells. (A) Flow cytometry quantification of mean fluorescence intensity (MFI) indicating IL-1R1 expression in the sLN cell populations. (B) Flow cytometric quantification of metastatic cells and (C) metastatic ratio in wild type and IL-1R1 knock-out mice, three weeks p.t.i.. (D) Proliferation assay (MTT) of B16-F1 treated with recombinant IL-1α for 24 h in comparison to untreated cells. (E) Proliferation assay (MTT) of human melanoma A375 treated with human recombinant IL-1α for 72 h, in comparison to untreated cells. (F) qPCR quantification of the Myd88 gene in B16-F1 stimulated with recombinant IL-1α in comparison to unstimulated.

### IL-1α promotes aggressiveness of melanoma metastasis via STAT3

To study the pathways influenced by IL-1α blocking *in vivo*, we performed scSeq of dissected metastases from mice treated with anti-IL-1α antibody at three weeks p.t.i. (Supp. Fig. 5A). Next, we performed an influence analysis on the transcriptomic scSeq data (Fig. 5A) to identify the top ten IL-1α key players, defined as the genes with the highest influence from all the *il1a* related pathways (Fig. 5B). Amongst them, we focused on *STAT3*, which was the most differentially expressed gene among the *il1a* key players following IL-1α depletion (Fig. 5C). This gene codifies for the transcription factor STAT3, a well characterized mediator of aggressiveness in different cancers, including melanoma^53,54^. Furthermore, scSeq analysis of STAT3 expression highlighted the metastatic melanoma as the cells exhibiting the highest expression levels of this transcription factor (Fig. 5D). Moreover, the induction of the *STAT3* gene in melanoma cells by IL-1α was confirmed *in vitro* by the administration of recombinant IL-1α (Fig. 5E). To recapitulate this mechanism at a functional level, we studied STAT3 protein expression and phosphorylation in murine melanoma cells using immunoblot assay. Treatment with recombinant IL-1α induced a significant overexpression of STAT3 in comparison to untreated controls starting at 24 h post IL-1α administration, while the addition of the anti-IL-1α depleting antibody reverted this phenotype (Fig. 5F). Furthermore, exposure to recombinant IL-1α induced the phosphorylation of STAT3, which was prevented by the depletion treatment (Fig. 5G). To evaluate if this mechanism was also present in a human model, we quantified STAT3 and pSTAT3 in the A375 cell line post administration of human recombinant IL-1α, and we observed a significant increase of both total and phosphorylated forms in comparison to untreated controls (Fig. 5H and 5I, respectively). Moreover, IHC sections of sLN from human patients confirmed the expression and phosphorylation of STAT3 in the metastatic lesions (Supp. Fig. 5B, C). Once we confirmed the connection between IL-1α and STAT3, we evaluated if a therapy with a STAT3 inhibitor (STAT3i) was able to improve the efficiency of the previously described IL-1α blocking therapy in the model of metastatic melanoma. Firstly, we observed that the administration of the combined therapies *in vivo* was able to contain the growth of the metastases more effectively than each of the two individual treatments (Fig. 5J). Additionally, we evaluated the combinatorial effect of these two therapies using an *in vitro* system, which highlighted a synergistic effect of the anti-IL-1α blocking therapy combined with the STAT3i stattic (Fig. 5K, Supp. Fig. 5D, E). In more detail, the addition of stattic improved both the efficacy (Supp. Fig. 5F) and the potency (Supp. Fig. 5G, H) of the IL-1α blocking therapy.

**Figure 5.**
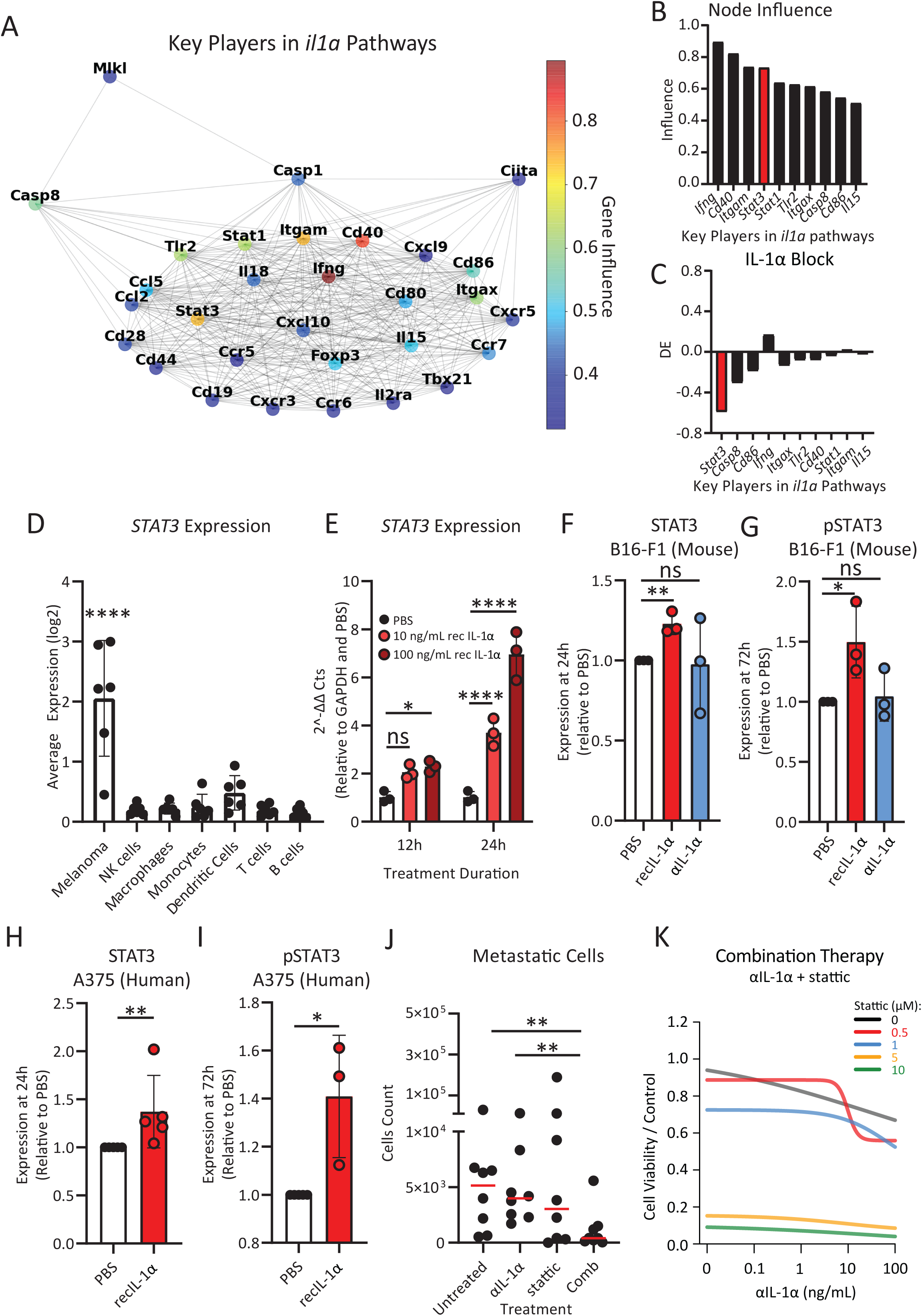
IL-1α induces STAT3 expression and phosphorylation in tumor. (A) STRING graph representing the most influential genes obtained by node influence analysis of il1a enriched pathways in scSeq data of the metastatic area of the sLN. The influence of each node is expressed in a colorimetric scale. (B) Bar plot showing node influence of the ten most influential genes in il1a pathways three weeks p.t.i.. STAT3 is highlighted (red). (C) Bar plot indicating differential expression (DE) of the ten most influential nodes in tumor following IL-1α block in comparison to untreated mice. STAT3 is highlighted (red). (D) Average STAT3 expression in each cell population of the metastatic sLN. (E) qPCR quantification of STAT3 expression in B16-F1 following recombinant IL-1α administration. Quantification of (F) STAT3 and (G) pSTAT3, measured by immunoblot, in B16-F1 after IL-1α treatment. Immunoblot quantifications of (H) STAT3 and (I) pSTAT3 in human melanoma A375 following exposure to IL-1α. (J) Flow cytometric quantification of metastatic cells in mouse sLN of mice treated with anti-IL-1α antibody, the STAT3 inhibitor stattic or their combination, in comparison to untreated. (K) Proliferation of mouse melanoma upon combination therapy with anti-IL-1α stattic at different concentrations, measured by MTT assay.

## DISCUSSION

In the present work, we characterized the innate immune responses of the sLN to melanoma metastasis invasion. We discovered that the SSM, which are the first immune cells to encounter melanoma metastasis in the sLN, phagocyted malignant cells and released IL-1α. Rather than triggering a tumor-killing inflammation, this cytokine increased metastatic cells aggressiveness by promoting STAT3 phosphorylation and increasing cancer proliferation. Importantly, blocking IL-1α decreased metastatic growth and cooperated synergistically with STAT3 inhibition.

STAT3 is a transcriptional factor with a relevant role in melanoma progression^55^, together with relevant immunosuppressive and pro-angiogenic properties^56,57^. Diverse studies investigated the STAT3 pathway and its activation by IL-6^58^. Of note, no studies reported similar effects of IL-1α. Based on this gap of knowledge, we stress a new target for metastatic melanoma therapy, acting on STAT3 signaling. This finding has further relevance in the context of combined therapies, which represent a very promising approach to target cancer cells at different levels, including tumor microenvironment^59,60^. In this context, blocking multiple immune pathways, such as IL-1α and IL-6, might improve the efficacy of STAT3i in comparison to single or dual therapy, as suggested by other studies indicating the synergistic effect of these two cytokines^36^. In addition, considering the variability of cytokines levels and responses to cytokines-based therapies in patients^61^, IL-1α blockade could be envisaged as an alternative to IL-6 inhibition for boosting STAT3i^62^ in those patients with low levels of IL-6 and low sensitivity to IL-6 depletion^63^. The specific cytokines expression profiling in patients, in fact, might be a useful tool to predict the patient response to the treatment and to design the best therapeutic strategy, according to the concept of personalized medicine, as previously proposed^64–66^. Moreover, IL-1α blocking agents have already been tested in clinical trials on patients with various tumors and with different grading, showing variable efficiency^41,42,44,64^. Moreover, previous evidence, described a possible connection between PD-1 and IL-1^38,67^ or STAT3^57,68^ in various tumors, indicating that IL-1α blockade in combination with other therapies, including checkpoint inhibitors, might be object of further studies.

Another important aspect of the present work is the focus on metastasization. Metastases are in fact more dangerous than primary tumors for patients, especially in melanoma^69^. Moreover, their biology might differ substantially from the primary tumor^70^. For instance, STAT3 favors the spread of melanoma cells to distant organs, and it is particularly expressed in the melanoma metastasis^53,54^. For this reason, the identification of the IL-1α – STAT3 axis, able to address efficiently metastases at their first stage, gains particular clinical relevance^48,71^. Furthermore, the pro-tumoral effect of SSM over the metastatic melanoma might be associated with the more aggressive phenotype acquired by these cells in comparison to the primary tumor or to peripheral blood circulating melanoma cells^11,12^. However, other mechanisms, such as, for instance, the lymph-mediated protection from ferroptosis^13^, are also involved in this process.

In this study, we clearly characterized the pro-tumoral activity of SSM during melanoma metastasis. However, previous evidence reported controversial functions of SSM in tumor biology^9,25,72^. These cells, indeed, belong to the family of macrophages, which are capable of activating both pro- and anti-tumoral mechanisms, as a consequence of their cellular plasticity^73^. For this reason, targeting a specific pathway, as we proposed here, might reveal a better strategy than depleting the whole macrophage population, avoiding the hampering of possible anti-metastatic functions of these cells^23,24,74^. Similarly, the development of drugs capable of targeting specifically SSM might reveal useful to block the IL-1α – STAT3 axis only in these cells and to boost their anti-tumoral properties, in a process of macrophages re-polarization^75^. Unfortunately, despite recent studies described compounds capable of localizing differentially in the SCS and in the medullary area of the LN, a therapy able to differentiate specifically macrophage subsets in the LN is still missing ^72,76,77^.

Additionally, SSM initiate the inflammatory response in the sLN by different mechanisms, including cell death associated with the release of pre-stored IL-1α^20,78^, which functions as an alarmin molecule following release from dying cells^79^. However, despite we observed a prominent recruitment of immune cells, we have not detected an efficient tumor-killing. Different hypotheses could explain this observation, including but not limiting to the exhaustion of the innate immunity, a higher affinity for IL-1α in melanoma cells in comparison to immune cells, or a specific inhibition of IL-1R1 by IL-1R antagonist in the immune compartment, which has been previously suggested in a study in human patients^80^. These and other hypotheses will furnish exciting insights on novel methods to improve immunotherapy and should be investigated in the future.

In conclusion, our results provide evidence of a novel function of SSM in melanoma metastasis progression by controlling the IL-1α – STAT3 axis. Importantly, IL-1α blocking decreased metastasis growth and acted synergistically with a STAT3i in controlling tumor growth. Taken together, these findings provide with new opportunities to improve currently available immunotherapies against metastatic melanoma.

## MATERIALS AND METHODS

### Cell culture and lentiviral transduction

B16-F1, B16-F10 and A375 cell lines were provided, respectively, by G. Guarda (IRB, Bellinzona) and C. Catapano (IOR, Bellinzona). E0771 cell lines were acquired from Ch3 BioSystems. All cell lines were maintained in a complete RPMI medium (RPMI+Hepes, 10 % heat inactivated FBS, 1 % Glutamax, 1 % sodium pyruvate, 1 % non-essential amino acids, 50 units/mL Penicillin, 50 μg/mL Streptomycin and 50 μM β-mercaptoethanol) and regularly tested for mycoplasma (MycoAlert Mycoplasma Detection kit, Lonza). The B16-F1-mCherry and B16-F1-Azurite cell lines were generated by lentiviral transduction. Briefly, lentiviral plasmids pSicoR-Ef1a-mCh (Addgene 31847) or pLV-Azurite (Addgene 36086) were transfected in HEK293T cells with packaging vectors pMD2G and psPAX (Addgene 12260 and 12259) to generate viral particles. After concentration by centrifugation, the virus was later collected and used for B16-F1 transduction. Transduced fluorescent cells were selected by live cells sorting using BD FACSAria Sorter.

### Mice

The Institute for Research in Biomedicine (IRB) hosted animal experiments in facilities defined as specific pathogen-free facilities, according to FELASA guidelines. Experiments involving IL-1α KO mice were conducted at the Ben Gurion University animal facility. In both facilities mice were housed in Individually Ventilated Cages (IVC) with controlled light: dark cycle (12 : 12), room temperature (20 - 24 °C) and relative humidity (30 - 70 %). Animal caretakers, researchers and veterinarians provided mice with daily check of general health conditions. All animal experiments have been conducted in accordance with the Swiss Federal Veterinary Service guidelines and the Israel Animal Welfare Act. All mouse procedures have been previously authorized by the relevant institutional committee (Commissione Cantonale per gli Esperimenti sugli Animali) of the Cantonal Veterinary Office and by the by the Israeli Council for Animal Experimentation of the Ministry of Health, with licensing numbers TI 25/2017, TI 24/2018, TI 55/2018 and TI 30/2020. Charles River Laboratories, F. Sallusto (IRB, Bellinzona) and R. Apte (BGU, Be’er Sheva) provided, respectively, C57BL/6J, B6.129S7-Il1r1tm1Imx/J (IL-1R1KO/KO, Jackson code 003245) and IL-1α KO mice^81^, which were then bred in-house. B6.129P2(Cg)-Cx3cr1tm1Litt/J (CX3CR1GFP/wt) mice were originally acquired from Jackson Laboratories (cat 005582) and bred in-house. The genotype of all mice was confirmed as previously described^82,83^. Mice in an age range of 6 - 12 weeks, showing good health conditions and no abnormal clinical signs took part in the experiments. Equal numbers of males and females were assigned to experimental groups through a statistical randomization process. Power calculation per groups size determination, performed by using R software (R: A Language and Environment for Statistical Computing, R Core Team, R Foundation for Statistical Computing, Vienna, Austria), estimated a number of 10 animals per group to obtain > 99 % statistical power.

### Allograft model

10^6^ cells from the syngeneic cell lines B16-F1, B16-F10 and E0771 were injected subcutaneously in the right footpad in 10 μL sterile PBS. Mice were anesthetized with Isoflurane (5 % for induction, 3 % for maintenance, FiO2 = 1, 1 L/min) and monitored, after cells injection and anesthesia recovery, to check for absence of pain or impaired movement. Mouse body weight and tumor size were measured every one or two days. Tumor volume was calculated with the formula V = (length x width^2^) / 2 and mice were euthanized when tumor reached 250 mm^3^. Euthanasia was performed by isoflurane overdose followed by cervical dislocation and immediate organs collection. We excluded from experiments mice which did not develop tumor (V = 0 mm^3^ at day 20 p.t.i.) or which developed tumor in the popliteal fossa, impeding the collection of the popliteal lymph node. In some experiments, we injected 15 μL of the B16-F1 tumor cell lysate originated from 5 × 10^5^ cells, sonicated at constant cycles of 30 seconds.

### *In vivo* treatments

To maximize the specific effect of treatments on LN metastases and to minimize the effect on tumor engraftment and primary tumor growth, all treatments were administered when the primary tumor reached a size of 40 mm^3^, which corresponds to the time of arrival of the first metastatic cells to the LN. Additionally, all local treatments were injected in the calf, to minimize their distribution to the primary tumor. All treatments were resuspended in a maximum volume of 10 μL in calcium- and magnesium-free PBS (PBS−). After injection in the subcutis, mice were recovered from anesthesia and monitored for absence of any sign of pain in the foot. Carrier-free recombinant mouse IL-1α (Biolegend) was locally administered at a dose of 1 μg / 10 μL per day. The anti-mouse IL-1α monoclonal antibody (InVivoMAb anti-mouse IL-1α, clone ALF-161, BioXCell) was administered to deplete IL-1α at a dose of 200 μg i.v. plus 60 μg locally, as previously reported^20^. Depletion was then maintained with a daily local injection of 60 μg. STAT3 was inhibited by local injection of stattic (SelleckChem) 3.75 mg/kg every two days. Stattic was reconstituted, according to manufacturer’s instructions, in 5 % DMSO (VWR), 40 % PEG300 (MedChem Express), 5 % Tween® 80 (Sigma-Aldrich) and 50 % distilled water. For macrophages depletion, mice received locally 10 μL of clodronate- or PBS-containing liposomes (Liposoma), followed by a second dose two days later.

### IVIS

To monitor primary tumor growth and mCherry expression of fluorescent cancer cells, we used the IVIS Spectrum Imaging System (Caliper LifeSciences). Mice were anesthetized with isoflurane as above described to measure epifluorescence. Immediately after image acquisition, animals were recovered from anesthesia. Images were later analyzed using Living Image Software 4.2 (Caliper LifeSciences).

### Immunofluorescence and immunohistochemistry

For mouse microscopy experiments, organs were fixed immediately after collection in 4 % paraformaldehyde (Merck-Millipore) for 12 h at 4 °C, then washed in calcium- and magnesium-free PBS (PBS−) and embedded in 4 % Low Gelling Temperature Agarose (Sigma-Aldrich). 50 μm sections were cut with a vibratome (VT1200S, Leica). Slices were stained in a blocking buffer composed of TritonX100 (VWR) 0.1-0.3 %, BSA 5% (VWR) and fluorescently labelled antibodies at proper concentration, all diluted in PBS supplemented with calcium and magnesium (PBS+). After 72 hours of incubation at 4 °C, samples were washed in 0.05 % Tween® 20 (Sigma-Aldrich), post-staining fixed with PFA 4 %, washed in PBS- and mounted on glass slides. Confocal images were acquired using a Leica TCS SP5 microscope with a 20 × 0.7 oil objective. To quantify the rate of invasion of melanoma in each sLN region, we first identified metastatic mass on the mCherry channel with an automatic Otsu threshold, after noise filtering with ImageJ Despeckle plugin and size filtering for regions bigger than 30 μm^2^. LN regions were manually identified based on CX3CR1 and CD21/35 morphology. Next, we quantified the total tumor area and the percentage of overlap of metastasis with each other LN region, respectively. Sample sizes were distributed as follows: n = 21, 7 and 11 for week one, two and three p.t.i., respectively. To quantify the expression of CD169^+^, the LN regions were manually identified as described above and the mean fluorescence intensity of each region was later calculated. To stain IL-1α in human lymph nodes, samples were stained using the BOND-III fully automated IHC/ISH stainer (Leica Biosystems) according to the manufacturer’s instructions. To stain STAT3 and pSTAT3, primary antibodies (mouse anti-Stat3, clone 124H6, and mouse anti-Phospho-Stat3, Tyr705, clone M9C6, Cell Signaling) were incubated overnight at 4°C and the MACH4 Universal HRP-Polymer Detection System (Biocare Medical) was applied according to the manufacturer’s protocol. 3D cell reconstruction was performed using Imaris Cell Imaging Software (Oxford Instruments).

### Flow Cytometry

LNs were collected, disrupted with tweezers, and enzymatically digested for 10 minutes at 37 °C. DNase I (0.28 mg/mL, VWR), dispase (1 U/mL, Corning) and collagenase P (0.5 mg/ml, Roche) were resuspended in calcium- and magnesium-free PBS (PBS−). Digestion was stopped using a solution of 2 mM EDTA (Sigma-Aldrich) and 2 % heat-inactivated filter-sterilized fetal calf serum (Thermo Fisher Scientific) diluted in PBS− (Sigma-Aldrich). Fc receptors were blocked (αCD16/32, BioLegend) followed by surface staining and analyzed by flow cytometry on an LSRFortessa or FACSymphony (BD Biosciences). For IL-1α detection, intracellular staining was performed with a dedicated kit (88/8824/00, eBioscience), following the manufacturer’s instructions. Data were analyzed using FlowJo software (FlowJo LLC). To measure cytokines and chemokines expression in the LN, LEGENDPlex assays (Mouse Proinflammatory Chemokine Panel and Mouse Inflammation Panel; BioLegend) were used. Briefly, pLNs were collected and carefully disrupted in 75 μL ice-cold phosphate buffer, minimizing cell rupture. The suspension was centrifuged at 100 rcf for 5 minutes and the supernatant was collected. 25 μL supernatant was used for cytokines and chemokines detection. Samples were analyzed by flow cytometry on an LSRFortessa or FACSymphony (BD Biosciences) and data were analyzed using LEGENDPlex software (BioLegend).

### Antibodies

The list of antibodies used to stain mouse samples includes anti-CD21/35 (CR1/CR2, clone 7E9, BioLegend), anti-podoplanin (clone eBio8.1.1, Invitrogen), anti-CD3 (clone 17A2, BioLegend), anti-B220 (CD45R, clone RA3-6B2, BioLegend), anti-Gr-1 (clone RB6-8C5, BioLegend), anti-NK1.1 (clone PK136, BioLegend), anti-MHC II (I-A/I-E, clone M5/114.15.2, BioLegend), anti-CD11b (clone M1/70, BioLegend), anti-CD11c (clone N418, BioLegend), anti-F4/80 (clone BM8, BioLegend), anti-CD169 (Siglec-1, clone 3D6.112), anti-IL-1R1 (clone FAB7712N, R&D Systems), anti-IL-1α (clone ALF-161, BioLegend; clone REA288, Miltenyi Biotec), anti-CD4 (clone RM4-5, BioLegend), anti-CD8a (clone 53-6.7, Invitrogen), anti-CD25 (clone PC61, BioLegend). Human samples were stained with anti-IL-1α (clone OTI2F8, Novus Biologicals), and anti-CD68 antibodies (clone PG-M1, Dako).

### Single-cell RNA-sequencing

Metastatic LN were obtained from four PBS injected mice, six tumor-bearing mice and four tumor-bearing mice treated with anti-IL-1α as described above. Metastases from tumor-bearing mice were microsurgically dissected using sterile micro-surgical tools. SS, IF and F regions were dissected in negative controls. Later, samples were disrupted into single cell suspension as described for flow cytometry, using sterile nuclease-free tools. Single cells were barcoded using the 10x Chromium Single Cell platform, and cDNA libraries were prepared according to the manufacturer’s protocol (Single Cell 3′ v3, 10x Genomics, USA). In brief, cell suspensions, reverse transcription master mix and partitioning oil were loaded on a single cell chip, then run on the Chromium Controller. Reverse Transcription was performed within the droplets at 53 °C for 45 minutes. cDNA was amplified for a 12 cycles total on a Biometra thermocycler. cDNA size selection was performed using SpriSelect beads (Beckman Coulter, USA) and a ratio of SpriSelect reagent volume to sample volume of 0.6. cDNA was analyzed on an Agilent Bioanalyzer High Sensitivity DNA chip for qualitative control purposes. cDNA was fragmented using the proprietary fragmentation enzyme blend for 5 minutes at 32 °C, followed by end repair and A-tailing at 65 °C for 30 minutes. cDNA was double-sided size selected using SpriSelect beads. Sequencing adaptors were ligated to the cDNA at 20 °C for 15 minutes. cDNA was amplified using a sample-specific index oligo as a primer, followed by another round of double-sided size selection using SpriSelect beads. Final libraries were analyzed on an Agilent Bioanalyzer High Sensitivity DNA chip for quality control. cDNA libraries were sequenced on a NextSeq500 Illumina platform aiming for 50,000 reads per cell. Base calls were converted to reads with the software Cell Ranger (10x Genomics; version 3.1)

### Quality control, processing, annotation, and differential gene expression analysis of single-cell RNA-sequencing data

We used the cellranger pipeline^84^ to generate gene expression count matrices from the raw data. For each sample, a gene-by-cell counts matrix was used to create a Seurat object using Seurat^85,86^. We filtered cell barcodes with < 500 UMIs and > 5 % mitochondrial contents. Each individual sample was then normalized by a factor of 10,000 and log transformed (NormalizeData). The top 2000 most variable genes were then identified within each sample using the FindVariableFeatures method. We then integrated the cells from all samples together using FindIntegrationAnchors and IntegrateData (2000 genes). The integrated gene expression matrix obtained by applying the filtering steps above was then used to perform principal component analysis (RunPCA), preliminary clustering analysis, including nearest neighbour graph (FindNeighbors) and unbiased clustering (FindClusters), and cell type annotation. Uniform Manifold Approximation and Projection (UMAP) was then used to visualise the integrated expression data. We identified gene expression markers for each cluster using FindAllMarkers from Seurat with default settings, including Wilcoxon test and Bonferroni p value correction^85,86^. Differential gene expression between specified clusters (or subclusters) was performed using FindMarkers (Wilcoxon rank sum test) with Benjamini-Hochberg false discovery rate (FDR) correction, average log fold change (logFC) and detection/expression percentage rate (pct). Genes were considered (significantly) differentially expressed if FDR < 0.05, logFC > 0.2 and pct > 20 % within the cells in a given group.

### Gene relevance analysis of single-cell RNA sequencing data

To determine gene relevance across the single cell RNA sequencing data, we used a network science approach. To study nodes relevance we applied Graph Theory rules^87,88^ using mathematical and social network analysis concepts. We restricted the analysis to protein-protein interactions and to the pathways in which the gene *il1a* is involved. Relevant pathways (Cytokine-cytokine receptor interaction, Necroptosis, Hematopoietic cell lineage, Type I diabetes mellitus, Pertussis, Leishmaniasis, Tuberculosis and Inflammatory bowel disease) and related were selected using the Kyoto Encyclopedia of Genes and Genomes^89^. Then we measured the expression of these genes in our scRNA-Seq dataset and we used their expression values as input of STRING^90^. The resulting graph was used for the network analysis. We implemented a previously described comprehensive algorithm for evaluating node influences in social networks^91^. This algorithm is based on three centrality measures: eigenvector centrality^92^, current flow betweenness centrality^93,94^ and reachability^95^. Eigenvector centrality computes the centrality for a node based on the centrality of its neighbors. Current-flow betweenness centrality starts from an electrical current model describing the spreading pattern, to which betweenness centrality, which uses shortest paths, is applied. Finally, reachability refers to the local reaching centrality of a node in a directed graph as the proportion of other nodes reachable from that node. In addition single cell CMP values are taken as weights of the nodes. Basing on these parameters, the algorithm ranked node influences by analyzing preference relations and performing random walk. In the first step a partial preference graph (PPG) is derived from the analysis of preference relation between every node pair for each measure. Later, the comprehensive preference graph (CPG) originated from the combination of preference relations and the three previously indicated measures. Finally, a random walk was executed on CPG to determine node effect. By applying this implementation to scSeq data, it was possible to obtain a list of genes related to *il1a* pathways, according to their importance in our dataset.

### Proliferation (MTT)

To evaluate tumor cells proliferation and response to treatments, B16-F1 and A375 cells were seeded in a 96-well plate. Carrier-free recombinant mouse (Biolegend) or human (SinoBiological) IL-1α were administered at 10 ng/mL and cells were incubated, respectively, for 24 hours or 72 hours. Anti-mouse IL-1α monoclonal antibody (InVivoMAb anti-mouse IL-1α, clone ALF-161, BioXCell) was administered at the indicated doses 24 hours before data collection. To inhibit STAT3, stattic (SelleckChem) was administered at the indicated dosages. Proliferation was assessed by MTT (Methylthiazolyldiphenyl-tetrazolium bromide) assay according to the manufacturer’s recommendations (Sigma). Absorbance (OD, 560 nm) was measured in a microplate reader (Cytation 5, BioTek). Sensitivity to single drug treatments was evaluated by IC50 (4-parameters calculation upon log-scaled doses), as previously reported^96^. The beneficial effect of the combinations versus the single agents was considered both as synergism according to the Chou-Talalay combination index^97^, as previously performed^96,98^, and as potency and efficacy according to the MuSyC algorithm^99^.

### qPCR

To measure the expression of *STAT3, Myd88* and *Gapdh* genes, the following sets of primers were designed: *STAT3* forward, 5’-CACAAATATTTTTGAGTCGGCGC-3’; *STAT3* reverse 5’-AAAGCCCCCGATGAGGTAATTC-3’; *Myd88* forward, 5’-CGGCAACTAGAACAGACAGACT-3’; *Myd88* reverse, 5’-GCAAACTTGGTCTGGAAGTCAC-3’; *Gapdh* forward, 5′-ACATCATCCCTGCATCCACT-3′; *Gapdh* reverse, 5′-AGATCCACGACGGACACATT-3’. To isolate RNA from cell culture, cells were disposed of in single-cell suspension in calcium- and magnesium-free PBS (PBS−). RNA was isolated using an RNAeasy Mini kit (QIAGEN). Two μg of cDNA were synthesized using a cDNA synthesis kit (Applied Biosystems) following the manufacturer’s recommendations. For the qPCR reaction, a SYBR Master Mix (Applied Biosystems) was used, and samples were run on a QuantStudioTM 3 Real-Time PCR System (Thermofisher). mRNA levels were expressed relative to GAPDH expression. The Pfaffl method^100^ was used to calculate the relative expression of the transcripts.

### Immunoblotting

To evaluate protein expression in tumor cells, B16-F1 and A375 cells were treated using carrier-free recombinant mouse (Biolegend) or human (SinoBiological) IL-1α, at 100 ng/mL. To block IL-1α, anti-mouse IL-1α monoclonal antibody (InVivoMAb anti-mouse IL-1α, clone ALF-161, BioXCell) was administered at the dose of 100 ng/mL. All treatments were administered either for 24 hours or 72 hours. Cells were harvested and lysed by boiling samples in 2x Laemmli sample buffer (BioRad), supplemented with β-mercaptoethanol (Merck), for 10 minutes. Lysates (30-50 µg) were resolved according to molecular weight by electrophoresis using Mini-PROTEAN TGX Precast gels 4-20 % gradient (BioRad). Next, proteins were blotted onto nitrocellulose membrane (BioRad) by electric transfer and the membranes were blocked in TBST (20 mM Tris-HCl [pH 7.5], 150 mM NaCl, 0.1 % Tween 20) with 5 % nonfat dry milk (BioRad) for one hour at room temperature. The following primary antibodies were used in TBST 5 % BSA buffer: mouse monoclonal, anti-Stat3 (clone 124H6, 9139, Cell Signaling Technology) and rabbit monoclonal, anti-p(Y705)Stat3 (9131, Cell Signaling Technology). Mouse monoclonal anti-GAPDH (clone FF26A/F9, CNIO) was used in TBST with 5 % nonfat dry milk. The secondary antibodies used were: ECL anti-mouse IgG horseradish peroxidase-linked species-specific whole antibody and ECL anti-rabbit IgG horseradish peroxidase-linked species-specific whole antibody (GE Healthcare). Membranes were treated with Westar ηC 2.0 chemiluminescent substrate (Cyanagen) and signals were detected using digital imaging with Fusion Solo (Vilber Lourmat).

### Statistical analyses

All raw data were analyzed, processed and presented using GraphPad Prism 8.2.1 (Graphpad Software, La Jolla, USA). First, we applied the Shapiro-Wilk normality test to analyze the distribution of data. Then we compared means among groups using One-Way ANOVA or Unpaired t test for data with normal distribution, and the non-parametric Kruskal-Wallis or Mann-Whitney test for groups which did not present a normal distribution. In all statistical tests P value is indicated as * when <0.05, ** when <0.005, *** when <0.0005, **** when < 0.0001.

## ACKNOWLEDGMENTS

We thank Federica Sallusto (IRB, Bellinzona, Switzerland), Carlo Catapano (IOR, Bellinzona, Switzerland) and Greta Guarda (IRB, Bellinzona, Switzerland) for providing, respectively, IL-1R1 KO mice, A375, B16-F1 and B16-F10 cell lines. We also thank David Jarossay (IRB, Bellinzona, Switzerland) for assistance in flow cytometry and live cell sorting experiments, Diego Ulisse Pizzagalli (IRB, Bellinzona, Switzerland) for the support in Imaris Software usage, Kevin Ceni (IRB, Bellinzona, Switzerland) for Supp. Mov. 1 editing, and Rocco D’Antuono (IRB, Bellinzona, Switzerland, now Francis Crick Institute, London, United Kingdom) for microscopy support.

