## Supplementary Figures for "IL-1α secreted by subcapsular sinus macrophages promotes melanoma metastasis in the sentinel lymph node by upregulating STAT3 signaling in the tumor"

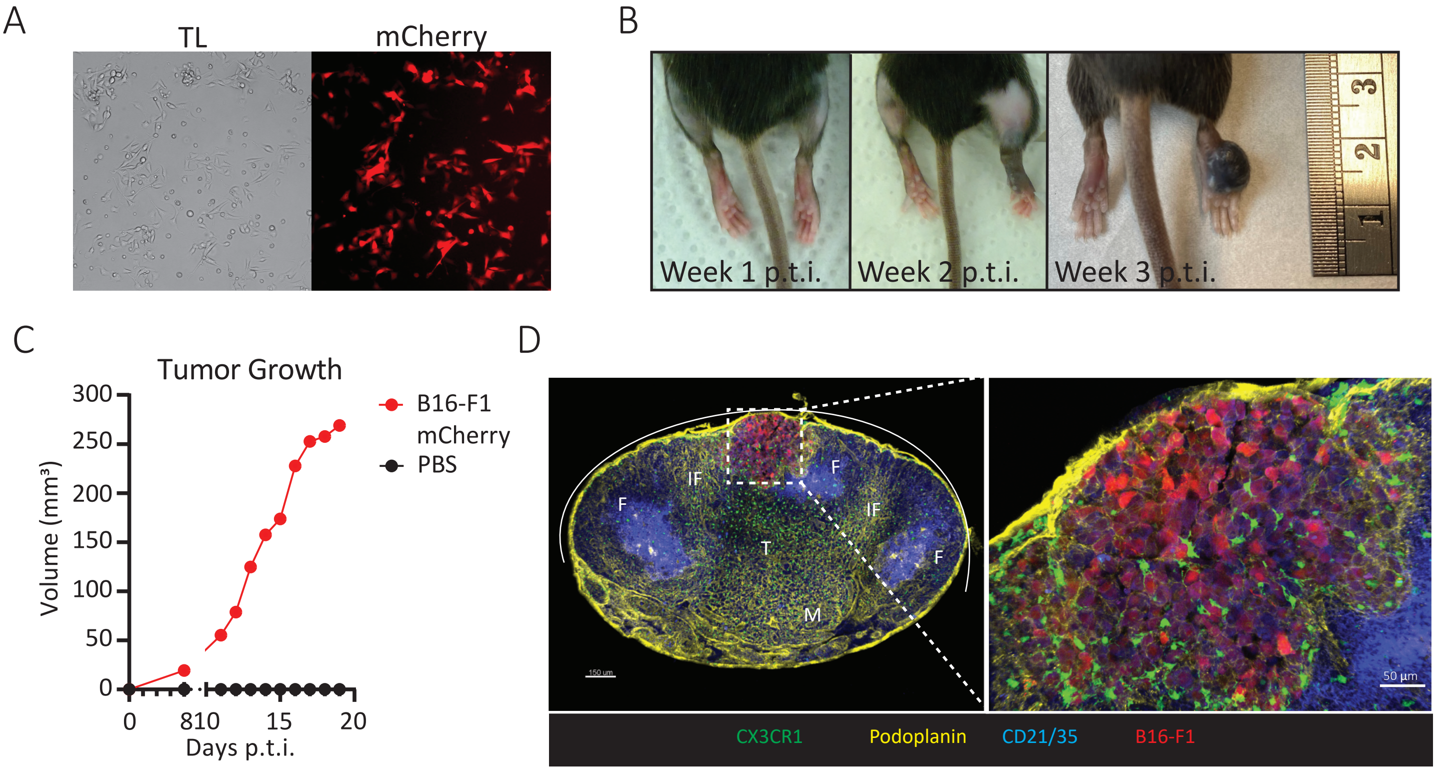


**Supplementary Figure 1 - Mouse melanoma metastases growth in the sentinel LN**

(A) Transmitted light (left) and mCherry (right) microscope pictures of transduced B16-F1 mCherry. (B) Representative pictures of primary tumor growth at week one, two and three p.t.i.. (C) Timecourse indicating tumor volume in the first three weeks p.t.i.. (D) Confocal micrograph showing sLN (left) at week three p.t.i. and magnification of the metastasis invading the SS and IF areas (right).

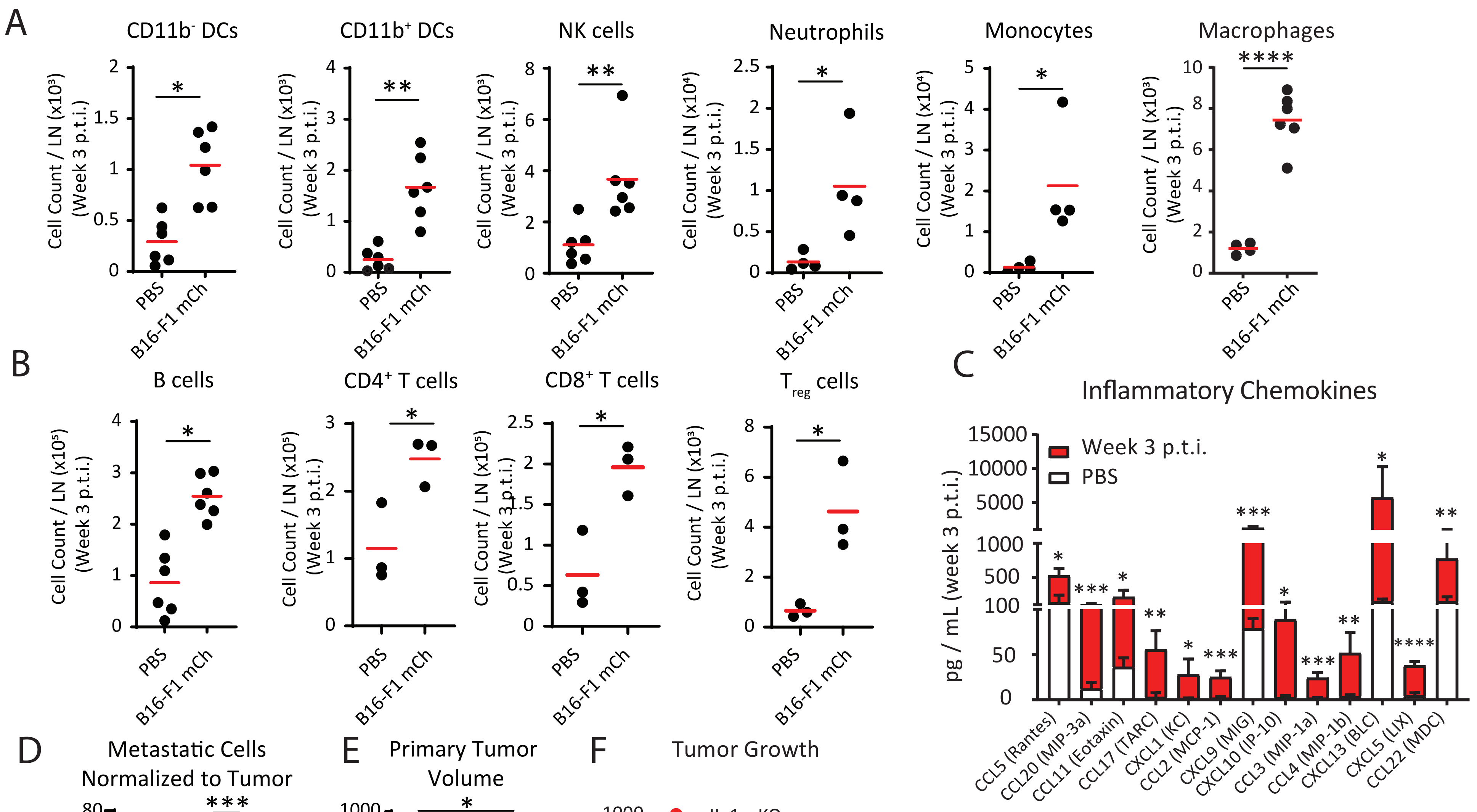


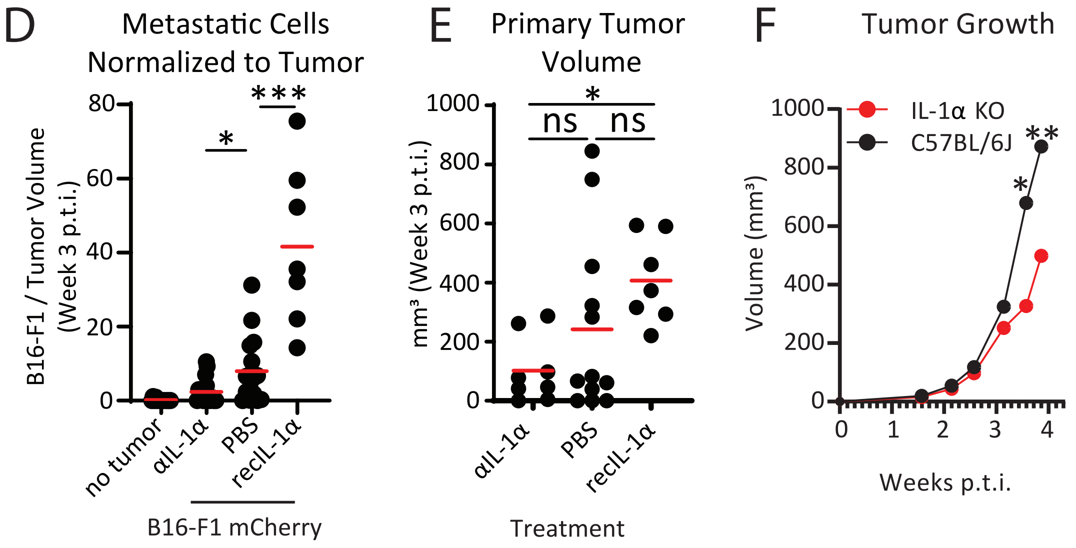


**Supplementary figure 2: Pro-tumoral release of IL-1α in the metastatic LN**
Total number of (A) innate and (B) adaptive immune cells in metastatic sLN at week three p.t.i.. (C) Inflammatory chemokines in metastatic sLN (red) in comparison to mice without tumor (white). (D) Number of metastatic melanoma cells in the sLN in mice treated with anti IL-1α or recombinant IL-1α, normalized to the primary tumor volume. (E) Primary tumor volume in mice treated with anti IL-1α blocking antibody or with recombinant IL-1α in comparison to untreated. (F) Primary tumor growth in IL-1α wild type and IL-1α KO mice.

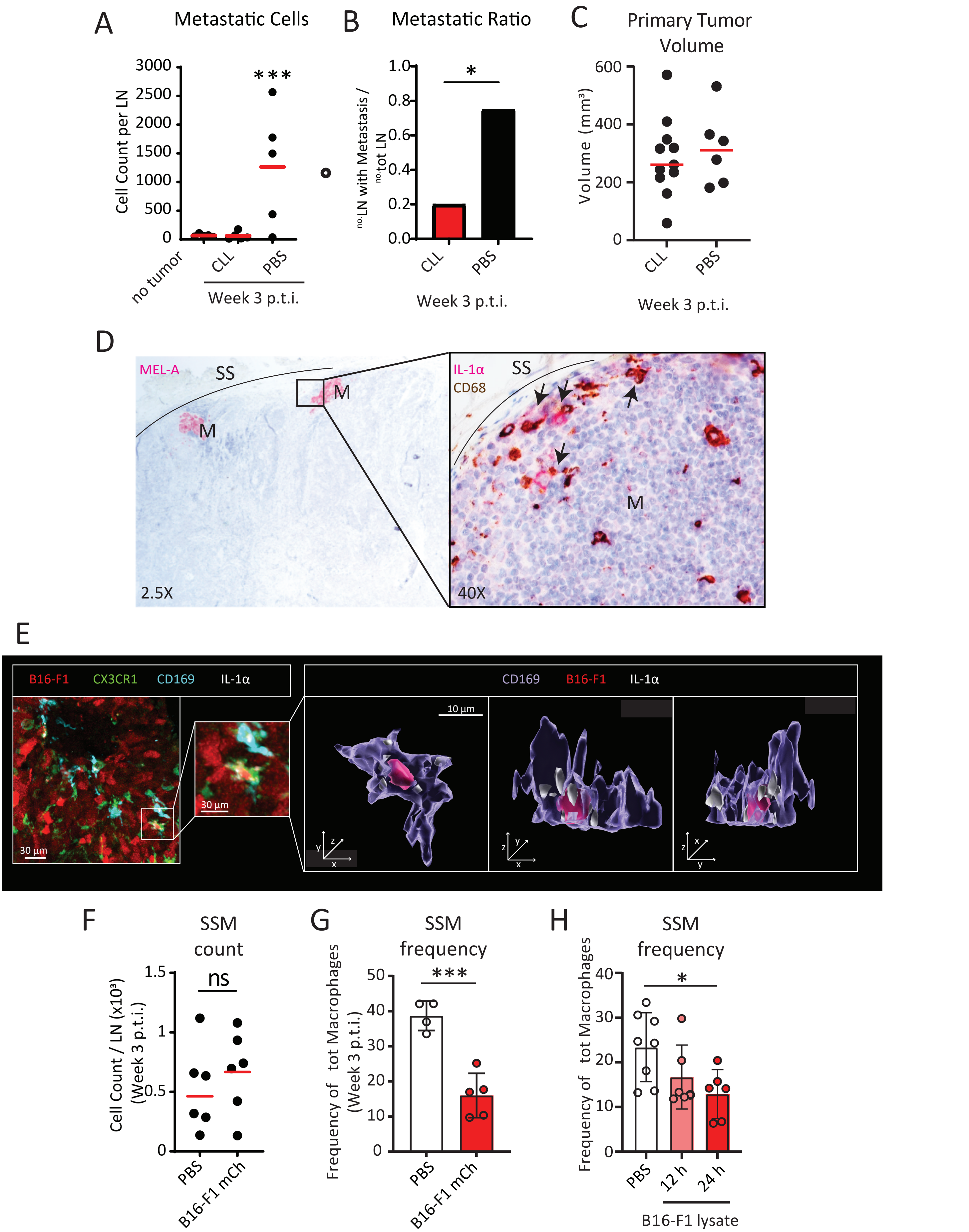


**Supplementary figure 3: SSM are the main source of IL-1α**

(A) Total number of metastatic cells in sLN of mice treated with clodronate liposomes (CLL) in comparison to negative controls. (B) Metastatic ratio and (C) Primary tumor volume in mice treated with CLL in comparison to untreated mice. (D, left) Representative section of human sLN indicating presence of melanoma metastases (pink), indicated with the letter M. (D, right) Magnification of metastatic area showing infiltration of CD68$^{+}$ macrophages (brown) and expression of IL-1α (pink). Representative macrophages positive to IL-1α are indicated by black arrows. (E, left) Confocal micrographs showing a tumor infiltrating SSM and (E, right) corresponding 3D reconstruction indicating intracellular accumulation of tumor (red) and IL-1α (white). (F) Total number of SSM and (G) frequency on total LN macrophages at three weeks p.t.i.. (H) Frequency of SSM at 12 and 24 hours after injection of tumor cell lysate.

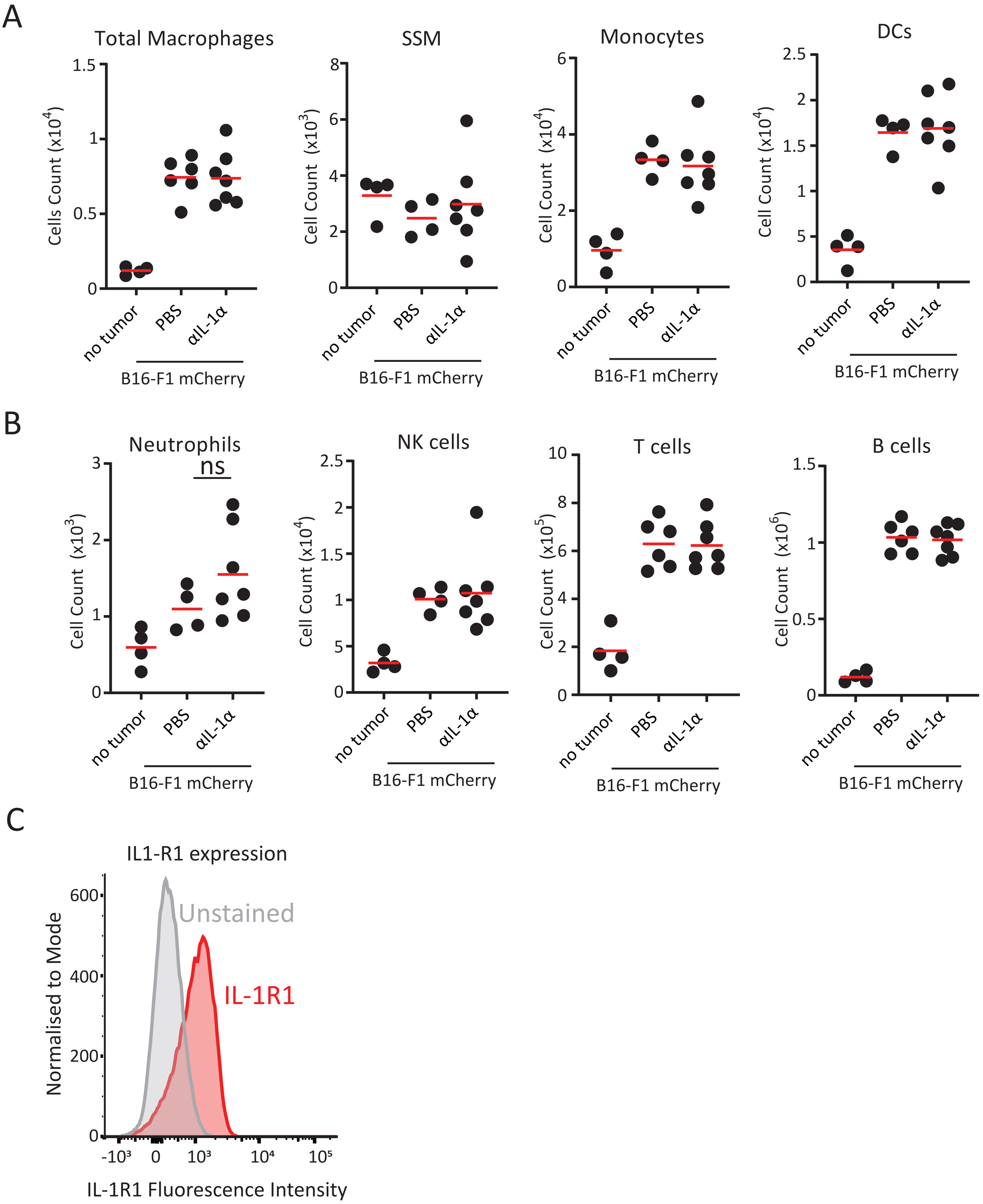


**Supplementary figure 4: SSM derived IL-1α directly supports tumor proliferation**
Total number of (A) phagocytic and (B) non phagocytic immune cells in sLN of mice treated with anti IL-1α antibody in comparison to untreated at three weeks p.t.i.. (C) Expression of IL-1R1 in B16-F1 *in vitro*.

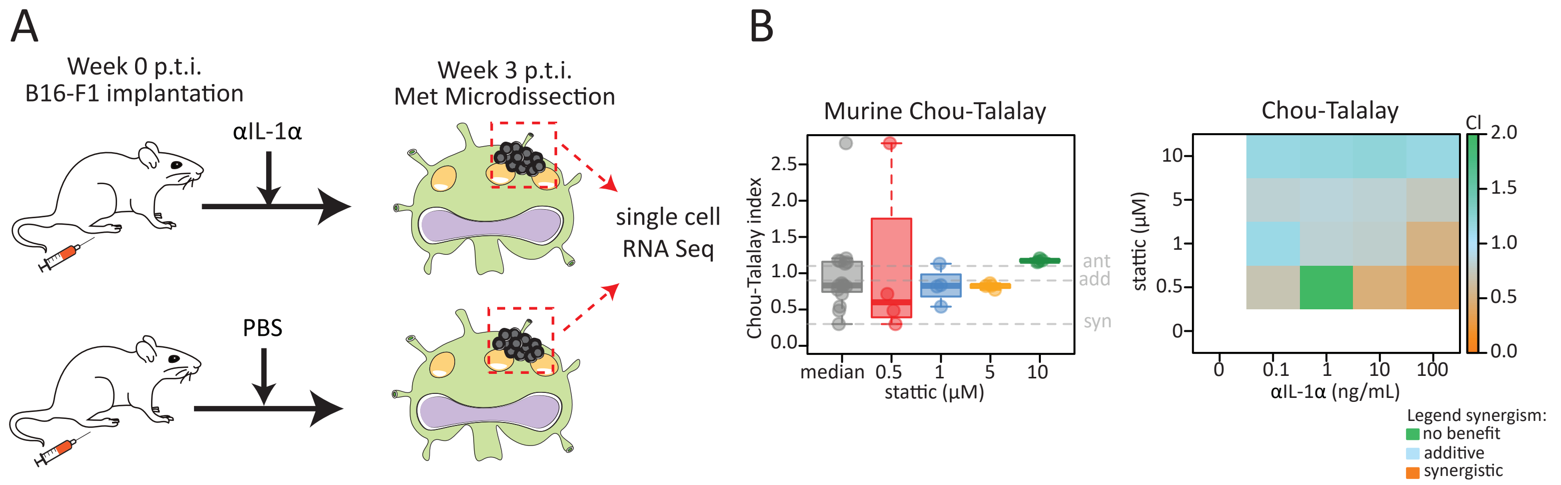


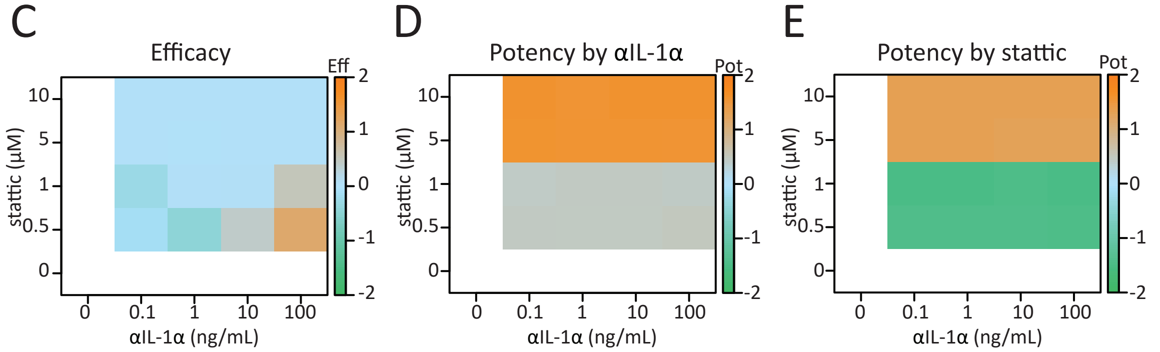


F

| Patiend ID | Clark grading | STAT3 (%) | pSTAT3 (%) |
|---|---|---|---|
| 1 | II | 10 | 20 |
| 2 | III | 80 | 15 |
| 3 | III | 60 | 60 |
| 4 | IV | 80 | 50 |
| 5 | IV | 50 | 5 |
| 6 | IV | <5 | <5 |
| 7 | IV | <5 | <5 |

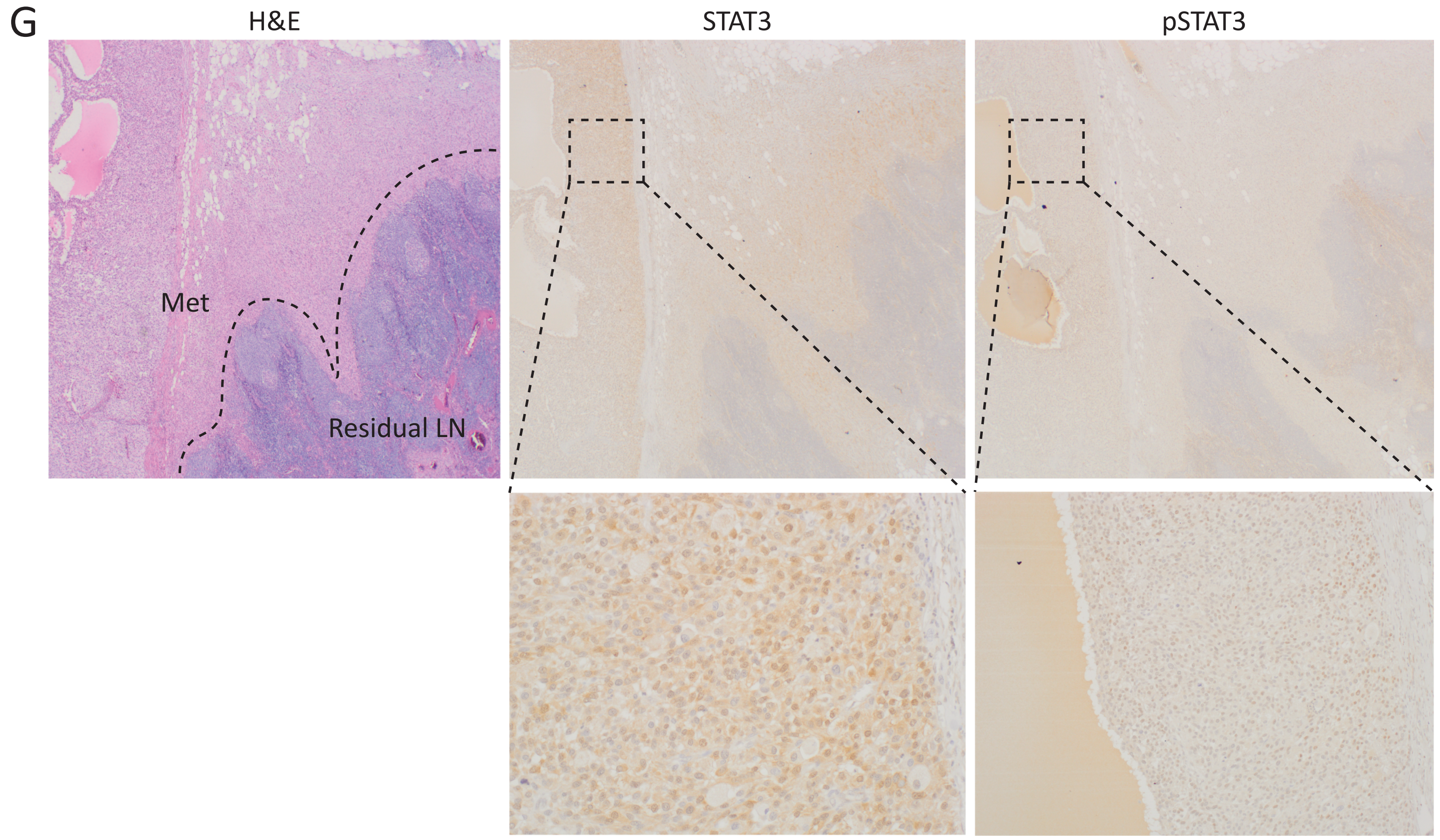


**Supplementary figure 5: IL-1α induces STAT3 expression and phosphorylation in tumor**
(A) Scheme representing the design of scSeq experiment. (B, left) Chou-Talalay Index distribution in samples, indicating the thresholds for synergistic, additive and antagonist effect in grey. (B, right) Chou-Talalay Index, (C) efficacy of the combination therapy, (D) potency by anti IL-1α and (E) potency by stattic, all indicated as average expression of samples. (F) Table reporting Clark grading and % of STAT3 and pSTAT3 positive cells in 7 human sLN metastatic samples. (G) Histological sections of a representative metastatic lesion in the sLN of a human patient, stained with haematoxylin and eosin (H&E, left) in comparison to IHC staining of STAT3 (center) and pSTAT3 (right), with corresponding magnifications (below).
